# Diverse *Escherichia coli* lineages, from domestic animals and humans in a household, carry colistin resistance gene *mcr-1* in Ecuador

**DOI:** 10.1101/350587

**Authors:** María Fernanda Loayza, Liseth Salinas, Fernando Villavicencio, Tamayo Rafael, Stephanie Salas, José Villacís, Carolina Satan, Liliana Ushiña, Ruth Rivera, Olga Muñoz, Jeannete Zurita, Tijet Nathalie, Roberto Melano, Jorge Reyes, Gabriel Trueba

**Author notes:** Address correspondence to María Fernanda Loayza.

## Abstract

The aim of this study was to investigate the presence of *Escherichia coli* carrying *mcr-1* gene in domestic animals close to a child who suffered a peritoneal infection by a *mcr-1* positive *E. coli*. Rectal or cloacal swabs and fecal samples from domestic animals were plated on selective media to isolate colistin-resistant *E. coli* and isolates were submitted to detection of *mcr-1* gene, pulsed field gel electrophoresis (PFGE), multi-locus sequence typing (MLST), replicon typing and S1-PFGE. Four *mcr-1* positive *E. coli* isolates (from chicken, turkey and dog) were recovered. No shared PFGE pattern or MLST sequence type were observed among isolates. A 60Kb IncI1**γ** *mcr*-1-carrying plasmid was detected in all isolates. Our results suggest that *mcr-1* gene was horizontally disseminated amongst different lineages of *E. coli* from domestic animals in the child’s household.

**Importance:** Horizontally transferable colistin resistance (*mcr*-1 gene) is thought to have originated in domestic animals and transferred to humans through meat and dairy products. In the present report we show evidence that the *mcr*-1 gene could be transferred to different *E. coli* strains colonizing different hosts (humans and pets) in the same household.

## Introduction

Domestic animals are important source of antibiotic resistant bacteria (and genes) which can be transmitted to humans through the food chain and direct contact (1). Plasmids and other mobile genetic elements (MGEs) are involved in the transmission of antimicrobial resistance among different bacterial genera colonizing different animal species (2, 3).

The first report of a colistin resistance (CR) gene carried by plasmids (*mcr*-1) came from China in 2015 (4). This gene codes for a phosphoethanolamine transferase (MCR-1), which modifies the lipid A moiety in the outer membrane of Gram negative bacteria and confers resistance to polymyxins (4, 5). Among Enterobacteriacea different *mcr* gen groups (1–5) could be transferred by mobile elements (6–8) and have been detected in humans, food animals, environmental samples in several countries, in different bacterial species and in several plasmid types (9). Enterobacteriacea has been found as the main *mcr* reservoir (6–8). The gene is easily transferred between commensal *E. coli* colonizing domestic animals and humans (7, 10) and opportunistic pathogens such as *E. coli* ST3941 (11) or *Klebsiella pneumoniae* ST512 KPC-3 (7, 12, 13).

In Ecuador, Ortega *et al.* have described the isolation of a colistin resistant *E. coli* carrying *mcr-1* from the peritoneal liquid in a child with complicated peritonitis (14). We investigated the *mcr-1* gene in *E. coli* isolates from domestic animals in this child’s household.

## Results

Four colistin resistant *E. coli* isolates were recovered from a dog (2 strains), a turkey (1 strain) and a chicken (1 strain). Susceptibility test for each strain in shown in **Table 1**. All CR isolates (colistin (MIC >4 μg/mL) were also resistant to ceftriaxone (MIC ≥64 μg/mL) and ciprofloxacin (MIC ≥4 μg/mL). Aditionally, dog isolates were resistant to ampicillin/sulbactam (MIC ≥32 μg/mL) and one of their isolates were resistant to gentamicin (MIC ≥16 μg/mL).

**Table 1.** Antibiotic susceptibility profiles and molecular analyses (PCR for *bla* genes detection, MLST and Replicon typing) of *mcr-1*-positive *E. coli* isolates.

| Strain origin | Antimicrobial resistance | MLST | PCR | Plasmid Inc group |  |  |  |  |  |  |  |  |  |  |
| --- | --- | --- | --- | --- | --- | --- | --- | --- | --- | --- | --- | --- | --- | --- |
|  |  | ST | *bla* genes | HI1 | I2 | N | FIA | FIB | I1y | FIIS | R | X1 | Y | FII |
| Chicken | COL, CRO, CIP | 3941 | *bla*<sub>CTX-M-65</sub> | - | + | - | + | - | + | - | - | - | - | - |
| Turkey | COL, CRO, CIP | 1630 | *bla*<sub>TEM-1</sub>; *bla*<sub>CTX-M-65</sub> | + | - | - | - | - | + | - | - | - | - | - |
| Dog1 | COL, SAM, CRO, CIP | 2170 | *bla*<sub>TEM-1</sub>; *bla*<sub>CTX-M-3</sub> | - | + | - | - | + | + | + | - | - | + | + |
| Dog2 | COL, SAM, CRO, CIP, GN | 2170 | *bla*<sub>TEM-1</sub>; *bla*<sub>CTX-M-3</sub> | - | - | - | - | - | + | - | - | - | - | - |
| Child* | COL, CAZ, CRO, FEP, CIP | 609 [13] | *bla*<sub>CTX-M-55</sub> [13] | - | + | + | - | - | + | - | + | + | - | - |

COL, colistin; SAM, ampicillin/sulbactam; CAZ, ceftazidime; CRO, ceftriaxone; FEP, cefepime; GN, gentamicin; CIP, ciprofloxacin.
ST, sequence type obtained by MLST analysis; *bla* genes, β-lactamase genes; Poultry TC, Transconjugant strain obtained with conjugation using
chicken CR *E. coli* isolate as donator and *E. coli* J53 as receptor strain. Replicon typing analysis was performed with PBRT KIT (DIATHEVA):
only positive Inc groups in the overall analysis are included. \* isolate reported by Ortega *et al*

Sequenced PCR products showed to be identical among them and close related to *mcr*-1 sequences that were previously reported (NCBI Accession number: KX11520.1, KX011521, KU935449.1, KU935446.1, KU935447, NG050417, KY013597).

Both PFGE and MLST indicated that CR *E. coli* isolates from domestic animals and the child (14) were different (Figure 1, Table 1). Sequence types (STs) carrying the *mcr-1* gene in the patient was ST609 and in the animal samples ST3941, 2170 and 1630. The isolates recovered from chicken and turkey also harbored bla_CTX-M-65_ (the one from turkey was blaTEM-1 as well), and the 2 *E. coli* from the dog were positive for bla_CTX-M-3_ and bla_TEM-1_ genes. All CR isolates from domestic animals in the household and the isolate from the child had an IncI1**γ** plasmid (Table 1). We were unable to transfer the *mcr*-1 gene using the conditions described in methods section. S1-PFGE showed a ≈ 60Kb plasmid positive for *mcr*-1 gene in all isolates (from animal and human) (Figure 1).

**Figure 1.**
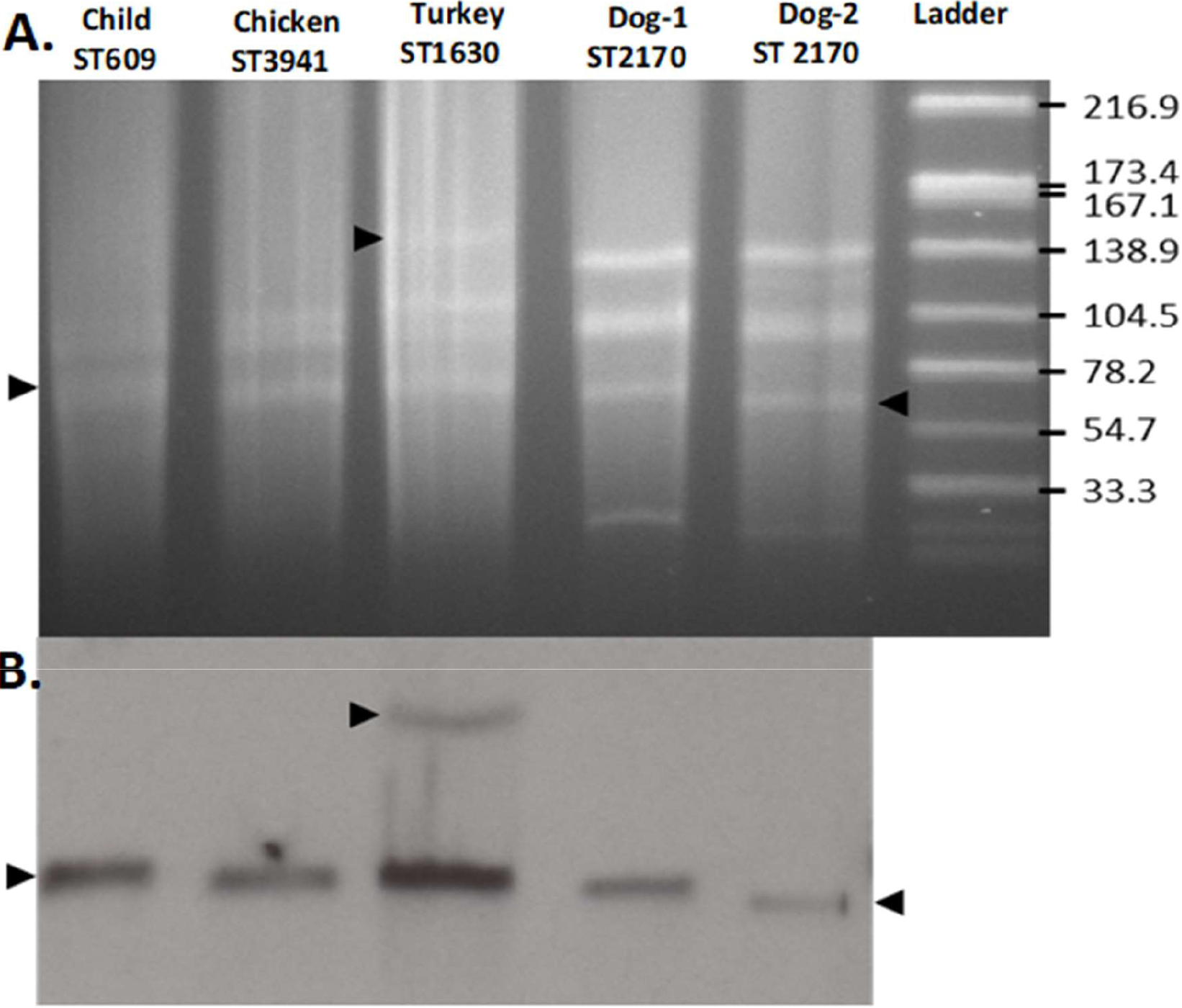
Identification of mcr-1-harboring plasmids. (A) S1- PFGE plasmid profiles of the patient and animals mcr-1 positive E. coli. (B) Southern blot using mcr-1 probe. Arrows indicate the plasmids carrying mcr-1 gene. Ladder, reference standard Salmonella enterica serotype Braenderup strain H9812 restricted with XbaI (sizes are given in kilobases)

## Discussion

*E. coli* genetic diversity has been studied in different environments showing no clear association patterns among different sources (15). In this study, colistin resistant *E. coli* strains isolated from different sources belonged to different lineages and no one was comparable with child isolate despite of they have an identical sequence.

*E. coli* ST 609 (child strain) has been previously detected in human (commensal and opportunistic pathogens), and water isolates in Spain, Argentina, United States, Vietnam, Netherlands, Australia, and Norway. ST 3941(chicken strain) has been identified in commensal bacteria from livestock and poultry in China and Vietnam farms, and *E. coli* ST 2170 (turkey strain) has been reported as an opportunistic pathogen causing septicemia in Japan and in healthy poultry farms in Vietnam. ST 1630 (Dog strains) was isolated from chickens in Denmark and United States (reported data in http://mlst.ucc.ie/mlst/dbs/Ecoli; last accessed December 8, 2017). Moreover, ST 1630 was also isolated from poultry in Japan (16). These data show the diverse and spread sources of *E. coli* strains detected and support the idea that successful mobile genetic elements could be responsible of antimicrobial resistance transference among dominant strains of commensal bacteria in different niches.

We were unable to transfer the *mcr*-1 gene to an acceptor, sodium azide resistant *E. coli.* Previous reports have described a low transfer frequency of *mcr*-1-carrying plasmids that could depends of plasmid type (16). However, we cannot discard that the conditions used in our study were not the optimal for their horizontal transfer.

Using a specific *mcr*-1 probe, one of the isolates (from turkey) showed two positive bands by Southern blot analysis: the common plasmid (~61 kb) and another one at ~140 kb. Li R *et al* (2016), have reported double plasmid carrying *mcr*-1 gene in commensal *E. coli* strains isolated from healthy pigs in China farms (17,18). The genetic analysis of those plasmids showed a composite transposon Tn*6330* (2 copies of IS*ApI1* flanking the *mcr-1* gene) that can form a circular intermediate which mediates the insertion of the *mcr-1* gene cassette into the IncHI2 plasmid. Further analysis is required to describe the *mcr-1* genetic environment and complete structure of plasmids from *E. coli* strains isolated in our study.

Both S1-PGFE and replicon typing suggest that the *mcr-1* gene was present in the same plasmid in all CR isolates (from domestic animals and human). It is probable that ≈ 60Kb plasmids were responsible for the transference of colistin resistance; previous reports have shown transmission of antibiotic resistance from bacteria in domestic animals to human bacteria is carried out by plasmids and not by antibiotic resistant clones (3).

Our study corroborates the notion that Enterobacteriacea colonizing intestines in domestic animals (including companion animals) could be reservoir of *mcr*-1 gene (2, 19–22). There was no evidence of colistin treatment in animals sampled, however, the presence of this type of antibiotic in animal feed and supplements in Ecuador prevents us from rule out antibiotic selection of CR clones.

## Materials and Methods

### 2.1 *E. coli strain isolation and MIC* determination

A cross – sectional study was conducted to detect commensal *E. coli* carrying *mcr-1* gene. Thirty-two fecal samples from soil and ten rectal or cloacal swabs were taken from rabbits (n= 2), guinea pigs (n= 2), dogs (n= 2) and chickens (n= 4). Fecal samples (n=32) from soil were placed in sterile reservoirs and swabs were placed in Tryptic Soy Broth (TSB, BD™) (19). Samples were transported to the Antimicrobial Resistance Laboratory in the Instituto Nacional de Investigatión en salud pública “Dr. Leopoldo Izquieta Perez”, Quito. Samples were plated on Mac Conkey Agar plates (MKL, BD™) supplemented with 2 μg/mL of colistin methansulfonate (RICHET®) (20). Identification and antimicrobial susceptibility profiles of the CR isolates were performed using the VITEK®2 compact (bioMérieux) with AST 272 card. Colistin minimal inhibitory concentration (MIC) was performed using SensititreTM (23).

### 2.2 Conjugation assay

Conjugation was performed using CR *E. coli* isolates as donor strains and *E. coli* J53 strain (sodium azide-resistant) as recipient (17). Trans-conjugant selection was performed in Trypticase ™ soy agar (Difco BD) with colistin (0.5μg/mL) and sodium azide (100 μg/mL) (24).

### 2.3 Molecular typing

PCR was performed to detect *mcr* gene (4); amplicons were sequence and aligned using *mcr-*1 (NG_055582.1), *mcr-*2 (NG_051171.1), *mcr*-3 (NG_056184.1), *mcr*-4(MG822665.1), *mcr*-5 (MG384740.1) accession numbers with Geneius software. Pulsed field gel electrophoresis (PFGE) (25) and multilocus sequence typing (MLST) was performed on seven housekeeping genes to define clonal relatedness (26). Replicon typing was performed using a commercial kit (PBRT KIT, DIATHE, Fano, Italy) (20, 27, 28). β-lactamase genes (*bla*_CTX-M-1_, *bla*_TEM_, *bla*_SHV_) were detected using primers previously described (29). Briefly, 12,5mL of GoTaq ® Green Master Mix (Promega, Madison, USA) were mixed with 1μL of upstream primer, 10μM, 1μL of downstream primer, 10μM, lμL of DNA template and 9,5 μL of Nuclease-free water to complete a 25μL of reaction volume. The reaction mix were amplified using 2-minute of initial denaturation at 94°C followed by 40 cycles of DNA denaturation at 94°C (40 sec), annealing at 60°C (40sec) and extension at 72°C (1min). The final elongation step at 72°C for 5 min. Amplicons were detected by electrophoresis in a 2% agarose gel. For complete amplification and sequencing of the detected resistance genes, primers and conditions previously described were used (30).

### Sl-PFGE and Southern blot

Plasmid content of each *E. coli* isolate and their estimated sizes were determined by S1 endonuclease-digested genomic DNA and PFGE (S1-PFGE). Briefly, genomic DNA agarose plugs of each isolate were partially digested with the endonuclease S1(31). DNA bands were separated by PFGE under previously described conditions (32). Plasmids were transferred and immobilized on a nylon membrane and identified by Southern blot analysis, using specific *mcr-1* digoxigenin-labeled probes (Roche Diagnostics).

## Conclusions

Our study suggests a polyclonal dissemination of *mcr-1* gene in *E. coli* domestic animals and humans in an Ecuadorian household. A 60Kb IncI1**γ** plasmid carried the *mcr-1* gene in this study, which may have been transferred among different strains colonizing different hosts. This study supports the current notion that mobile gene elements are more important than bacterial clones in the transmission of antibiotic resistance genes from the microbiota in domestic animals to human microbiota (33).

Our results highlight the importance of controlling the use of antibiotics in domestic animals and possibly the need for more studies of AR in isolates and mobile genetic elements in them.

## Funding

This study was funded by The Instituto Nacional de Investigación en Salud Pública “Dr.Leopoldo Izquieta Perez”, Quito – Ecuador and Instituto de Microbiología, Universidad San Francisco de Quito

## Ethical approval

No required

## Competing interest

The authors declare that they don’t have any conflict of interests.

